# Monocrotaline treatment of the rat predisposes to altered lipopolysaccharide-induced lung responses

**DOI:** 10.64898/2025.12.21.695829

**Authors:** D. DeBellis, T. Penn, E. Cowell, J. M. Roach, L.P. Kris, J. McEvoy-May, S. Bihari, D-L Dixon, J. M. Carr

## Abstract

There is a need to understand pathogen driven lung disease and the rat is a laboratory model, widely used to study acute lung injury (ALI). Here the monocrotaline (MCT) rat has been investigated as a model of an inflammatory lung with developing pulmonary hypertension (PH) on which an ALI is superimposed.

14 days following a single systemic dose of MCT, the lung is functionally normal, but stimulation with lipopolysaccharide (LPS) results in an altered response. The MCT/LPS lung is morphologically similar to LPS alone, with respiratory mechanics showing increased elastance, reduced compliance and increased tissue resistance. Bronchioalveolar lavage (BAL) fluid demonstrated a cellular infiltrate, with large macrophage-like cells, increased secreted angiotensin converting enzyme-2 (ACE2) and total protein. The lung transcriptome is pushed towards a pro-inflammatory M1 phenotype, (interferon (IFN)-ϒ, interleukin 6 (IL6), CD68 and CD80) compared to LPS alone. Additionally, the MCT/LPS lung has gene transcription signatures for enhanced cell death and DNA damage responses, higher levels of multiple complement components, dysregulation of the renin-angiotensin pathway with reduced ACE2 and increased AGTR1, increased factors such as Erythroferrone that would increase iron levels, and increased fibrinogen and A2M, that would promote thrombosis.

Thus, the MCT-treated rat represents an animal with no overt clinical distinction but following LPS stimulation elicits an exacerbated pathogenic lung response. The MCT-treated rat is a simple model that might be beneficial for understanding an M1 lung pathology with activated complement, low ACE2, high iron, and a propensity to clot in the context of developing cardiovascular disease.

## INTRODUCTION

Acute respiratory distress syndrome (ARDS) can result from conditions such as sepsis and pathogen infections of the lung with no single universal ARDS pathophysiology (Ashbaugh et al. 1967; Laffey and Kavanagh 2017; Kaku et al. 2020). This was also apparent during the SARS-CoV-2 pandemic where the COVID-19 lung represents a diverse pathology (Bos et al. 2021; Lascarrou 2021). ARDS is associated with an inflammatory lung response, dysregulation of homeostatic systems such as coagulation, and changes in the lung such as immune cell infiltration, microthrombi and endothelial cell activation leading to oedema and fibrosis (Force et al. 2012). Laboratory rodents have long been utilised as models for ARDS, in particular larger rodents such as the rat, where multiple respiratory measures are logistically more feasible (Bates 2017; Irvin and Bates 2003) and stimuli such as lipopolysaccharide (LPS) (Dixon, De Smet, and Bersten 2009; Domscheit et al. 2020), bleomycin (Kadam and Schnitzer 2023) and oleic acid provide an acute lung injury (ALI) (Idell et al. 1989; Matute-Bello, Frevert, and Martin 2008). A 2020 review of experts defined experimental ALI with 4 ‘domains’ - histologically demonstrated tissue injury, alteration of the alveolar-capillary barrier, inflammation, and lung dysfunction – but challenged the time frame of ‘acute’ (Kulkarni et al. 2022). The original concept of an injurious response within 24 hours (hr) was suggested to extend to encompass up to 7-10 days, depending on the stimulus, with a goal to more closely reflect clinical ARDS (Kulkarni et al. 2022). For instance, ARDS in the COVID-19 lung most often presents 10-14 days after SARS-CoV-2 infection and coincident with the decline in viremia (Polak et al. 2020). The recognition that models of actue lung injury do not reflect the full complexity of clinical ARDS, including the evolving nature of disease over a more extended time (Laffey and Kavanagh 2017) is an important caveat of preclinical models.

Our laboratory has recently reviewed the changes in the lung of the monocrotaline (MCT)-treated rat; a historically well-defined and often utilised laboratory model of pulmonary hypertension (PH) (Gomez-Arroyo et al. 2012; Hill, Gillespie, and McMurtry 2017), alongside the COVID-19 lung and as a potential model of ALI and ARDS (Kris et al. 2025). In the MCT rat model, inflammatory and thrombotic changes in the lung are reported 7-14 days following a systemic, single dose of MCT (Gillespie et al. 1985; Lai, Olson, and Gillespie 1991; Wilson et al. 1992). These changes in the lung are concurrent with evolving cardiovascular changes, with clear PH emerging later around 28 days post-treatment. Right ventricle failure is a contributor to mortality in the MCT-treated rat with wall thickening and insufficiency developing early and prior to end stage PH (Holda et al. 2020). Right sided heart dysfunction is also emerging as a contributor to the complexity of ARDS (Yogeswaran et al. 2025).The study here aimed to test experimentally if a single systemic dose of MCT could stimulate an aberrant host response in the rat lung that would be consistent with an ALI model, and further may subsequently modulate the lung response to a second pathogenic insult, such as LPS.

Outcomes demonstrate that while MCT alone has no major impact on measurable lung morphology or function, the transcriptional and cellular environment of the lung of the MCT-treated rat is primed to profoundly respond in a different way to a subsequent LPS stimulus. This response is biased toward a M1-like macrophage-driven proinflammatory environment, with the transcriptome suggesting heightened involvement of damaging complement pathways, in the context of increased cell death and dysregulation of iron homeostasis and the renin-angiotensin system (RAS). This model may be useful to study a more slowly developing lung and cardiovascular environment in an aged rat that is aligned with worsened responses to pathogens. Similar biological changes may underpin the phenotype of patient lungs that are more susceptible to develop ARDS or COVID-19 after an acute pathogen exposure.

## MATERIALS AND METHODS

### Animals and MCT treatment

Male Sprague Dawley Rats (300-400g, *n*=30 total) were administered MCT (Merck, C2401) or 0.03% DMSO vehicle control by intraperitoneal injection. For MCT at 60 mg/kg, this was administered in two separate injections: 40 mg/kg intraperitoneal (IP) and 20 mg/kg sub-cutaneous (S-C), due to ethical restrictions for injection of a maximum volume of 1 ml/kg. Groups included control DMSO (n=4) or MCT (n=4). Rats were anaesthetised and 200 µl of low dose, 3mg/kg LPS delivered intratracheally, as previously (Dixon, De Smet, and Bersten 2009; Green et al. 2024) to rats that had been DMSO (n=4, LPS alone) or 14-day MCT treated (n=5, MCT/LPS). Use of animals was reviewed and approved by the Flinders University Animal Ethics Committee, approval number AEC BIOMED5696.

### Respiratory mechanics

At the end point of experiments, rats were anaesthetised with 3% inhaled isoflurane (Zoetis Australia) at 1.5-2 mL/min O2 for five minutes and respiratory mechanics measured as previously described (Bihari et al. 2024). In brief, intraperitoneal injections of 50 mg/kg Thiopentone Sodium (Lyppard Australia Pty Ltd) were administered every five minutes, gradually reducing isoflurane from 1% to 0%. A tracheostomy was performed, and lungs were ventilated via a small animal mechanical ventilator (flexiVent; SCIREQ Scientific Respiratory Equipment, Montreal, Canada) at respiration rate – 120 breaths/min, tidal volume – 7 ml/kg, positive end-expiratory pressure (PEEP) – 3 cm/H2O. Prior to the measurement of lung mechanics, spontaneous breathing was prevented by paralysis through intravenous injection of vecuronium bromide (2 mg/kg), and a single 6 second sigh breath of 2.5x tidal volume (20 ml/kg) and 2 cm/H2O PEEP, applied to normalise lung volume history. This was immediately followed by a forced oscillation of 16 seconds of interrupted ventilation consisting of 19 sine waves with multiple prime frequencies (0.25 to 19.125 Hz) and a peak-to-peak volume excursion of 1.0 mL above the end-expiratory lung volume. The data were fitted to the constant phase model, with parameters as previously described (Bihari et al. 2024; Dixon et al. 2016).

### Tissue Collection

Following respiratory mechanics measurements the trachea, lungs and heart were resected *en bloc*. The lungs were lavaged three times with 3 volumes of cold 0.9% sodium chloride at 32 ml/kg to collect bronchoalveolar lavage (BAL) fluid. BAL cells were collected by centrifugation and counted by light microscopy. The upper right lobe of the lung was analysed for wet lung weight then freeze dried for dry lung weight. The middle and bottom right lung lobes were snap frozen for RNA analysis. The left lung lobe was gravity fixed in 10% (v/v) formalin for histological analysis.

### Histological staining

Lung tissue was paraffin embedded and 5µm sections were stained with hematoxylin and eosin (H&E). Lung morphology was scored by an independent blinded investigator, as previously described (dos Santos et al. 2011). 1×10^6^ BAL cells were fixed in 4% paraformaldehyde and DiffQuik modified Giemsa staining (Australian Biostain) was performed. Images were captured by light microscopy (Olympus BX53).

### ACE2 ELISA

ACE2 levels were determined by sandwich ELISA (Novus Biologicals, Cat: NBP2-78735) and total proteins in BAL fluid were determined via BCA Protein Assay (Thermo Fisher Scientific) as per manufacturer instructions.

### RNA extraction and RT-PCR

Methodology is as described previously (Green et al. 2024). In brief, total RNA was extracted with TRIzol (Invitrogen, Thermo Fisher Scientific), and 500 ng of DNase I-treated RNA reverse transcribed and RT-qPCR was performed using PowerUp SYBR Green Master Mix (Applied Biosystems, Thermo Fisher Scientific) with 1 µM of each primer as described in supplemental Table 1. Real time RT-qPCR was performed with a hot start protocol using a Rotor-Gene real-time PCR system (Qiagen). Results were normalised against the housekeeping gene, GAPDH and relative mRNA abundance determined by the ΔCt method.

### RNAseq and data analysis

Total RNA was extracted was extracted using the PureLink RNA Mini Kit (Invitrogen), assessed via Qubit and RNA libraries were prepared and sequenced on 2 MGI DNBSEQ G400 FCL (PE100) lanes by the South Australian Genomics Centre. Reads were aligned to reference genome GRCr8 and read quality control was performed through the NF-Core RNAseq (v3.22.0) pipeline under default parameters (Ewels et al. 2020; Patel 2025), with trimming through FastQC alignment with STAR and quantification with Salmon (Dobin et al. 2013; Patro et al. 2017). The NF-Core pipeline differentialabundance (v1.5.0) was used under default parameters with a false discovery rate cutoff of 0.05 to obtain differentially expressed genes (DEG) through the R package DEseq2 (Love, Huber, and Anders 2014). R package was also used to generate heatmaps (pheatmap), volcano plots (EnhancedVolcano with a log2FC threshold of ±1.5) and Gene ontology and KEGG pathway analyses (ClusterProfiler) (Yu et al. 2012).

### Data analysis

Data were collated in excel and graphs generated in GraphPad Prism10. Normally distributed data were analysed by Ordinary One-way ANOVA with Tukey’s multiple comparisons test.

## RESULTS

### MCT treatment for up to 14 days has no major impact on lung function, lung or liver morphology

Aged male rats, (approximately 12 months), were treated with 15mg/kg MCT or vehicle (DMSO) and respiratory mechanics analysed at 3- or 7-days post treatment or 60 mg/kg with analysis at 14 days post treatment. MCT treatment had no significant impact on respiratory mechanics, lung wet weight or cell counts in the BAL. Following H&E staining, morphologically, the lungs appeared normal (Supplemental Figure 1). The impact of 60 mg/kg MCT treatment on the liver at 14 days post treatment was assessed. No significant changes in mRNA for liver-derived complement components factor B (FB) and C3, and CXCL10 as a marker of inflammation were observed (Supplemental Figure 2).

### MCT-pretreatment alters the lung function, morphology and responses that are induced by LPS

Experiments next focused on 60 mg/kg MCT treatment with analysis at 14-day post treatment. The hypothesis was tested that although the lung physiology of the animal is not severely disrupted, MCT treatment will predispose the lung to an altered response to pathogen stimuli. Previously, lung stimulus of low dose (3mg/kg) LPS directly introduced into the lungs, induced minimal respiratory impact up to 6 h post administration (Dixon, De Smet, and Bersten 2009), and based on this our studies analysed 3mg/kg LPS, with or without prior MCT treatment. A single dose of 60 mg/kg MCT had no major impact on respiratory mechanics 14 days later (Figure 1A). In contrast, 24 hr LPS stimulation alone significantly increased elastance (Figure 1A). Pretreatment with MCT 14 days prior to LPS stimulation exacerbated adverse respiratory mechanics, with a significant increase in elastance and reciprocally a significant decrease in compliance accompanied by an increase in tissue dampening and resistance (Figure 1A). Gross analysis of the lung demonstrated a significant increase in wet and dry weight in MCT/LPS compared to DMSO or MCT alone, but no significant change compared to the lung treated with LPS alone (Figure 1B). Lung morphology demonstrated no gross morphological change with MCT alone although an increased cellular infiltrate in the alveolar spaces was present in both LPS and MCT/LPS treated lung tissue (Figure 1C). Pathology scoring of the lung for alveolar thickening and cellular infiltrate demonstrated no difference between DMSO or MCT treated lung tissue but a significant increase in both of these parameters in LPS and MCT/LPS treated lungs (Figure 1D).

**Figure 1.**
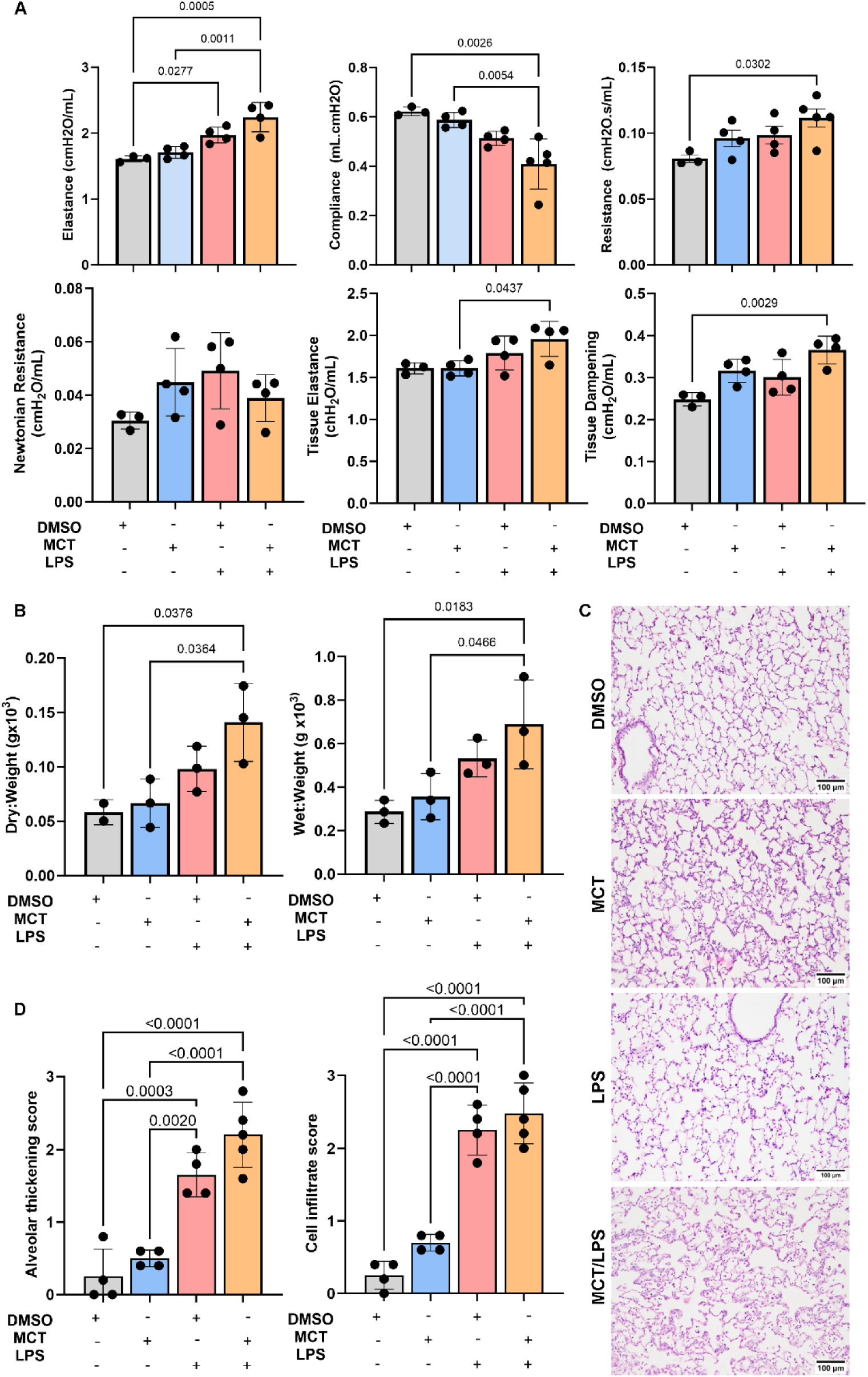
Prior MCT treatment exacerbates the adverse respiratory mechanics does not worsen the gross morphological changes in the lung induced by LPS treatment. Rats were treated with DMSO or 60mg/kg MCT and at 14 days post treatment vehicle or LPS delivered intratracheally. 24 h later **A.** respiratory mechanics were measured; and **B.** lungs weighed (wet) or following desiccation (dry); **C.** lungs were fixed and processed for H&E staining and images collected by brightfield microscopy. Representative images are shown; **D.** Images were scored blind by 2 independent investigators for alveolar thickening and cell infiltrate. Groups reflect DMSO alone (n=3); MCT alone (n=4); LPS alone (n=4) and MCT/LPS (n=5). Individual data points represent individual animals and mean ± SEM are shown. Data were analysed by one-way ANOVA, with Tukeys multiple comparison test and significance at p<0.05.

Consistent with a visual cellular infiltrate in the lungs seen by H &E staining, cell counts in the BAL fluid were significantly increased in LPS and MCT/LPS treated animals compared to DMSO alone (Figure 2A). To assess the cellular composition, 10^6^ cells from the BAL fluid was analysed were DiffQuik Giemsa stained (Figure 2B). The cellular composition of the BAL from DMSO or MCT-treated lungs contained mainly macrophage-like cells (Figure 2B). In contrast, the BAL from LPS or MCT/LPS lungs, contained a predominance of lymphocytic cells, with large macrophage-like cells in the MCT/LPS BAL (Figure 2B), with a higher magnification shown demonstrating large macrophage-like cells with prominent cytoplasm evident in the MCT/LPS rat BAL (Figure 2C). MCT treatment did not impact ACE2 levels in the BAL fluid, with a trend towards increased ACE2 with LPS alone, and significantly higher ACE2 in the BAL induced by combined MCT/LPS treatment (Figure 2D). This was associated with significantly elevated BAL total protein content from both LPS and MCT/LPS treated animals (Figure 2D).

**Figure 2.**
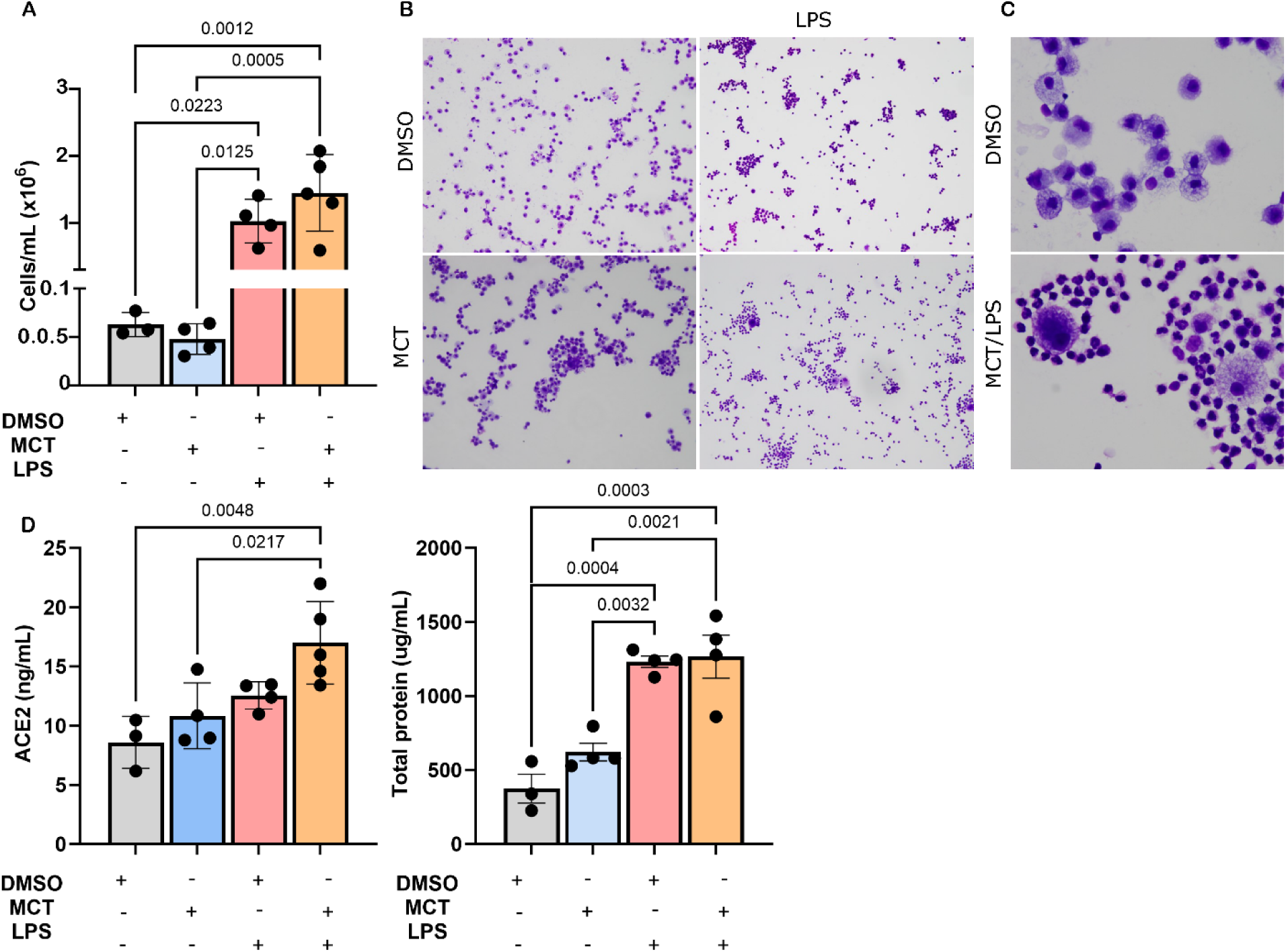
Increased cell number and altered morphology accompanied by increased ACE2 and total protein in the BAL of MCT/LPS treated rats. Rats were treated with DMSO or 60mg/kgMCT and at 14 days post treatment vehicle or LPS delivered intratracheally. 24 h later lungs were collected and BAL harvested. **A.** cells enumerated by brightfield microscopy. Results represent mean ± SEM, n=4 per group, n=5 MCT/LPS; **B.** BAL cells were resuspended at 10^6^/ml adhered to glass slides and Diffquick stained. Images were collected under bright field microscopy at 10X magnification; **C.** 40X magnification. Representative images are shown; **D.** ACE2 protein quantitated by ELISA, and total protein measured. Data were analysed by One-way ANOVA with Tukeys multiple comparisons, p values for significant difference are shown

The characteristics of the lung responses for inflammatory: IFN-ϒ, IL6, CCL5, TNF, CXCL10, C3; antiviral: viperin; or receptors for SARS-CoV-2: ACE2, TMPRSS2, wer analysed by RT-PCR. No significant induction of any target genes was observed following MCT treatment alone (Figure 3). LPS stimulation induced an inflammatory response with a significant increase in TNF, CXCL10 and C3, but a decrease in TMPRSS2 (Figure 3). The combination of MCT/LPS treatment did not dramatically impact on these LPS induced changes, except for a significant and clear increase in IFN-ϒ in MCT/LPS treated samples that was not seen with LPS or MCT alone (Figure 3).

**Figure 3.**
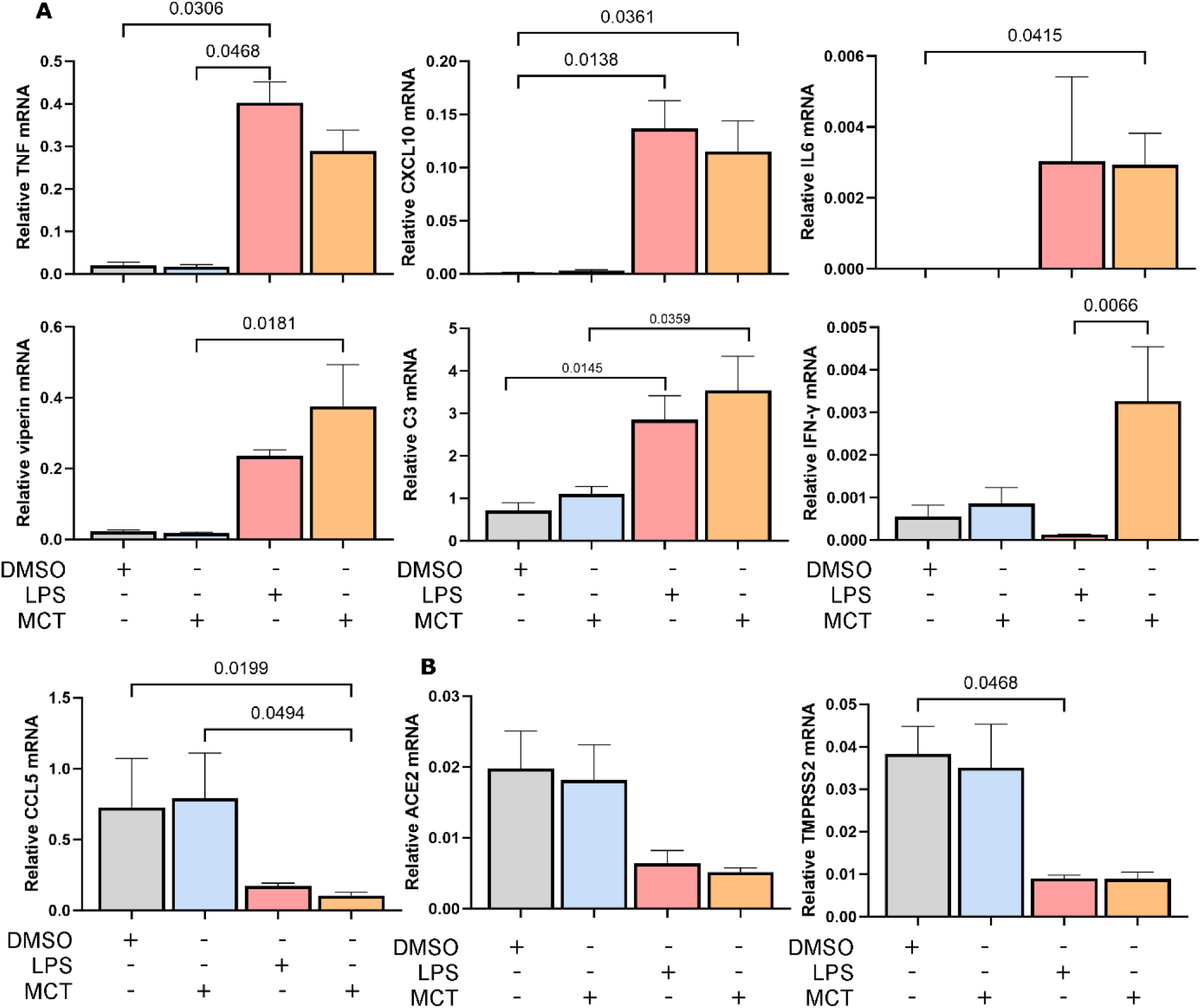
Inflammatory mRNA are induced in the LPS and MCT/LPS rat, with a unique induction of IFN-ϒ in MCT/LPS treated lung. Rats were treated with DMSO or 60mg/kg MCT and at 14 days post treatment vehicle or LPS delivered intratracheally. 24 h later lungs were collected, total RNA extracted and subjected to RT-PCR for **A.** inflammatory and antiviral factors; **B.** SARS-CoV-2 receptors. RT-PCR values were normalised to GAPDH and represent mean SEM, n=4 per group. Data were analysed by One-way ANOVA with Tukeys multiple comparisons, p values for significant difference are shown.

Overall, data suggests reduced lung respiratory function, morphological change to the lung tissue and BAL fluid, and a change in the characteristics of the cellular and cytokine/chemokine inflammatory response to LPS when rats were pre-treated with MCT.

### MCT impacts the lung transcriptome and alters LPS responses

Next, RNAseq was undertaken to define changes in the overall transcriptional profile of the lung response to LPS and/or MCT. Principle component analysis demonstrated separation of DMSO and two out of four MCT-treated animals, and a clear separation of animals treated with LPS and MCT/LPS (Figure 4A). Differential gene expression (DGE) suggests mRNA for 6 genes are upregulated uniquely by MCT alone, with one induced in both MCT and LPS alone treated animals, 166 in MCT and MCT/LPS and 64 mRNAs common to MCT, LPS and MCT/LPS (Figure 4A). LPS simulation induced 1625 unique mRNAs that were not induced by other treatments and 4275 mRNA common to both LPS and MCT/LPS (Figure 4A). MCT/LPS treatment induced 1570 mRNAs that were not observed in either MCT or LPS alone (Figure 4A). mRNAs in each portion of the Venn diagram are listed in the supplemental data file. These differences in gene expression were supported by heat map visualisation with two of the four MCT treated rats with distinct mRNA expression profile compared to DMSO alone, and distinct clustering of LPS and MCT/LPS treated animals (Figure 4B). Volcano plots demonstrate a small group of mRNAs upregulated in MCT-treated rats, with approximately 2-fold increase compared to DMSO (Figure 4C). In contrast LPS stimulated animals demonstrated significant up and down-regulated genes with much greater log-fold changes, and with a similar profile to MCT/LPS stimulated animals (Figure 4C). The comparison of MCT/LPS treated animals to LPS alone demonstrated a small group of significantly upregulated genes, with very few significantly downregulated genes (Figure 4C).

**Figure 4.**
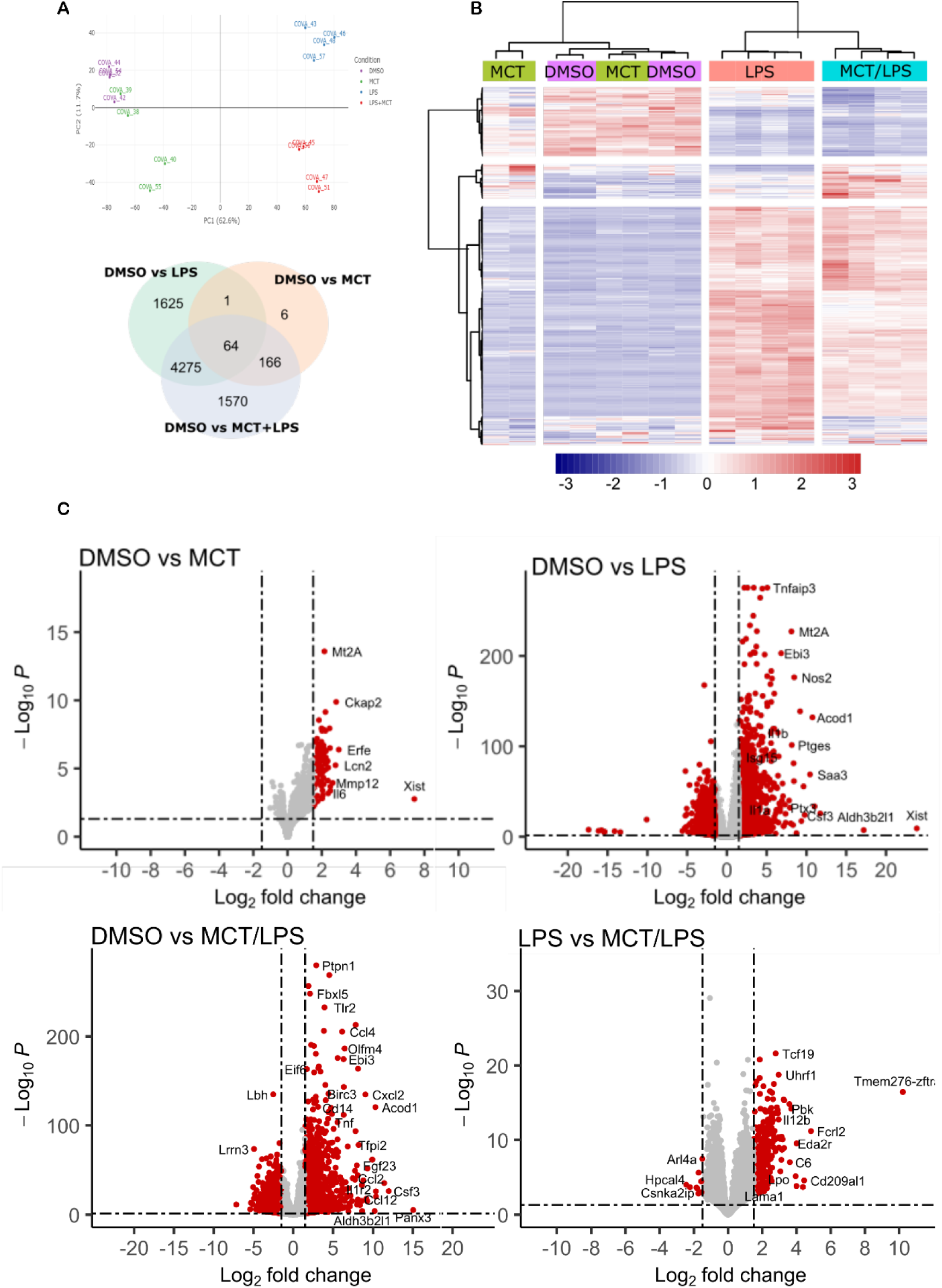
The transcriptome is different in MCT, LPS and MCT/LPS treated rats. Total RNA was extracted from rat lung tissue (n=4 per group) and subjected to RNAseq. **A.** Principle component analysis (PCA) of the top 500 most variable genes was generated with DEseq2. Venn diagram representation of the overlap of differentially expressed genes between treatment groups; **B.** Heat map representation of differentially expressed genes; **C.** Volcano plots of differentially expressed genes with a cut off value of 1.5 Log2 fold change.

Analysis of the top 20 up-regulated mRNAs in the MCT treated lung compared to DMSO, demonstrates upregulation of genes involved in iron homeostasis such as Erythroferrone (Erfe) and Lipocalin 2 (Lcn2), inflammation such as IL6, serum amyloid A3 (Saa3), and matrix metalloproteinase 12 (MMP12), and indicators of DNA damage (Dtl1, Ddias) (Table 1). Additionally, the top 20 mRNA changes for LPS and MCT/LPS compared to DMSO are summarised (Table 1). Comparison indicates induction of inflammatory molecules such as IL6, CXCL11, CSF3, and Nos2 and pentraxin-3 (Ptx3) in both LPS and MCT/LPS, with similar fold changes. Pannexin 3 (PanX3) was the highest fold change in the MCT/LPS treated lung (Table 1), that in comparison was only upregulated 1.4-fold in LPS alone. Aldh3b2l1 demonstrated a greater fold change in LPS compared to MCT/LPS (Table 1). Xist, a long non-coding RNA (lncRNA) involved in silencing the X chromosome but also associated with PAH in males (Carman et al. 2024), was the top fold change RNA in LPS stimulated animals and considerably lower at 2.7-fold in MCT/LPS. The top 20 mRNA changes in MCT/LPS compared with LPS alone indicated upregulation of mRNAs involved in inflammation such as the rat orthologue of CD209/DC-SIGN, paralemmin-3 (Palm3) a factor that is involved in LPS induced responses in alveolar macrophages (Chen et al. 2017), Osteopontin (Spp1) (Lin et al. 2023), complement component 6 (C6), and upregulation of MMP12, Dtl1, Ddias, as also seen with MCT alone (Table 1 and 2). A full list of mRNA changes is available in the associated supplemental data file.

**Table 1.** Top 20 differentially expressed genes in the MCT treated rat lung relative to DMSO and compared in LPS treated rats with the same genes seen in MCT/LPS treated rats. (# = P < number). Shaded items represent the equivalent gene on each list that is outside of the top 20 with distinct difference in log2fold change.

|  | Gene | log2FC | padj |  | Gene | log2FC | Padj |
| --- | --- | --- | --- | --- | --- | --- | --- |
| **1** | Xist | 7.412165 | 0.001734 | **11** | Il6 | 2.375031 | 0.000665 |
| **2** | Erfe | 2.993909 | 4.15E-07 | **12** | Saa3 | 2.356487 | 7.69E-05 |
| **3** | Ckap2 | 2.841348 | 1.31E-10 | **13** | Ccne1 | 2.347647 | 2.84E-06 |
| **4** | Lcn2 | 2.804667 | 5.78E-06 | **14** | Dtl | 2.267504 | 3.60E-06 |
| **5** | Mmp12 | 2.633003 | 0.000113 | **15** | E2f7 | 2.240975 | 5.31E-07 |
| **6** | Tnfrsf22 | 2.500066 | 0.000209 | **16** | Rmi2 | 2.238922 | 1.57E-06 |
| **7** | Ckap2-ps3 | 2.496899 | 0.000433 | **17** | Mybl1 | 2.22605 | 4.59E-06 |
| **8** | Pbk | 2.478921 | 3.19E-07 | **18** | Ddias | 2.219872 | 1.23E-05 |
| **9** | Ccne2 | 2.454875 | 1.14E-08 | **19** | Rrm2 | 2.2164 | 1.71E-08 |
| **10** | Xkr5 | 2.380124 | 8.68E-06 | **20** | Uhrf1 | 2.213187 | 7.22E-10 |
| **DMSO vs LPS** |  |  |  | **DMSO vs MCT/LPS** |  |  |  |
| **1** | Xist | 23.87556 | 7.19E-10 | **415** | Xist | 2.651981 | 0.00282 |
| **553** | Panx3 | 2.381634 | 0.003386 | **1** | Panx3 | 15.04467 | 7.00E-06 |
| **2** | Aldh3b2l1 | 17.19655 | 5.37E-08 | **15** | Aldh3b2l1 | 8.672767 | 8.98E-05 |
| **3** | Csf3 | 11.72909 | 2.15E-26 | **2** | Csf3 | 11.96244 | 3.25E-27 |
| **4** | Il6 | 11.01354 | 3.03E-34 | **3** | Il6 | 11.39587 | 3.04E-36 |
| **5** | Acod1 | 10.75869 | ##### | **6** | Acod1 | 10.28708 | ##### |
| **6** | Saa3 | 10.44319 | 1.44E-69 | **8** | Saa3 | 9.917357 | 1.27E-62 |
| **7** | Cxcl11 | 9.77823 | 1.01E-24 | **5** | Cxcl11 | 10.34912 | 5.52E-27 |
| **8** | Gpr84 | 9.635446 | 2.42E-56 | **10** | Gpr84 | 9.334354 | 1.04E-52 |
| **9** | Orm1 | 9.364424 | 5.23E-18 | **4** | Orm1 | 10.36821 | 5.54E-21 |
| **10** | Cxcl2 | 9.199844 | ##### | **13** | Cxcl2 | 9.077116 | ##### |
| **11** | Clca4l | 8.733563 | 0.000137 | **7** | Clca4l | 10.19511 | 7.97E-05 |
| **12** | Nos2 | 8.458821 | 0 | **20** | Nos2 | 8.148935 | ##### |
| **13** | Fgf23 | 8.426752 | 3.70E-62 | **25** | Fgf23 | 8.000212 | 3.74E-56 |
| **14** | Tfpi2 | 8.383164 | 6.92E-82 | **19** | Tfpi2 | 8.229727 | 9.66E-79 |
| **15** | Il24 | 8.309537 | 1.26E-13 | **11** | Il24 | 9.296868 | 7.41E-16 |
| **16** | Ptges | 8.154475 | ##### | **27** | Ptges | 7.833224 | 3.02E-94 |
| **17** | Mt2A | 8.106533 | ##### | **26** | Mt2A | 7.851964 | # |
| **18** | Cxcl6 | 8.029135 | 2.24E-15 | **12** | Cxcl6 | 9.193792 | 1.23E-18 |
| **19** | Ptx3 | 7.91613 | 9.32E-33 | **14** | Ptx3 | 8.820385 | 2.57E-39 |
| **20** | Cxcl10 | 7.821052 | 1.03E-40 | **28** | Cxcl10 | 7.772872 | 4.80E-40 |
| **22** | Ccl7 | 7.36486 | 5.15E-26 | **16** | Ccl7 | 8.636068 | 6.11E-34 |
| 25 | Il17c | 7.125897 | 2.23E-12 | 17 | Il17c | 8.622205 | 3.49E-16 |
| 77 | Serpina3a | 5.327681 | 9.78E-07 | 18 | Serpina3a | 8.394114 | 1.95E-11 |
| 52 | Ccl12 | 5.815309 | 6.88E-11 | 19 | Ccl12 | 8.132451 | 8.76E-18 |

**Table 2.**
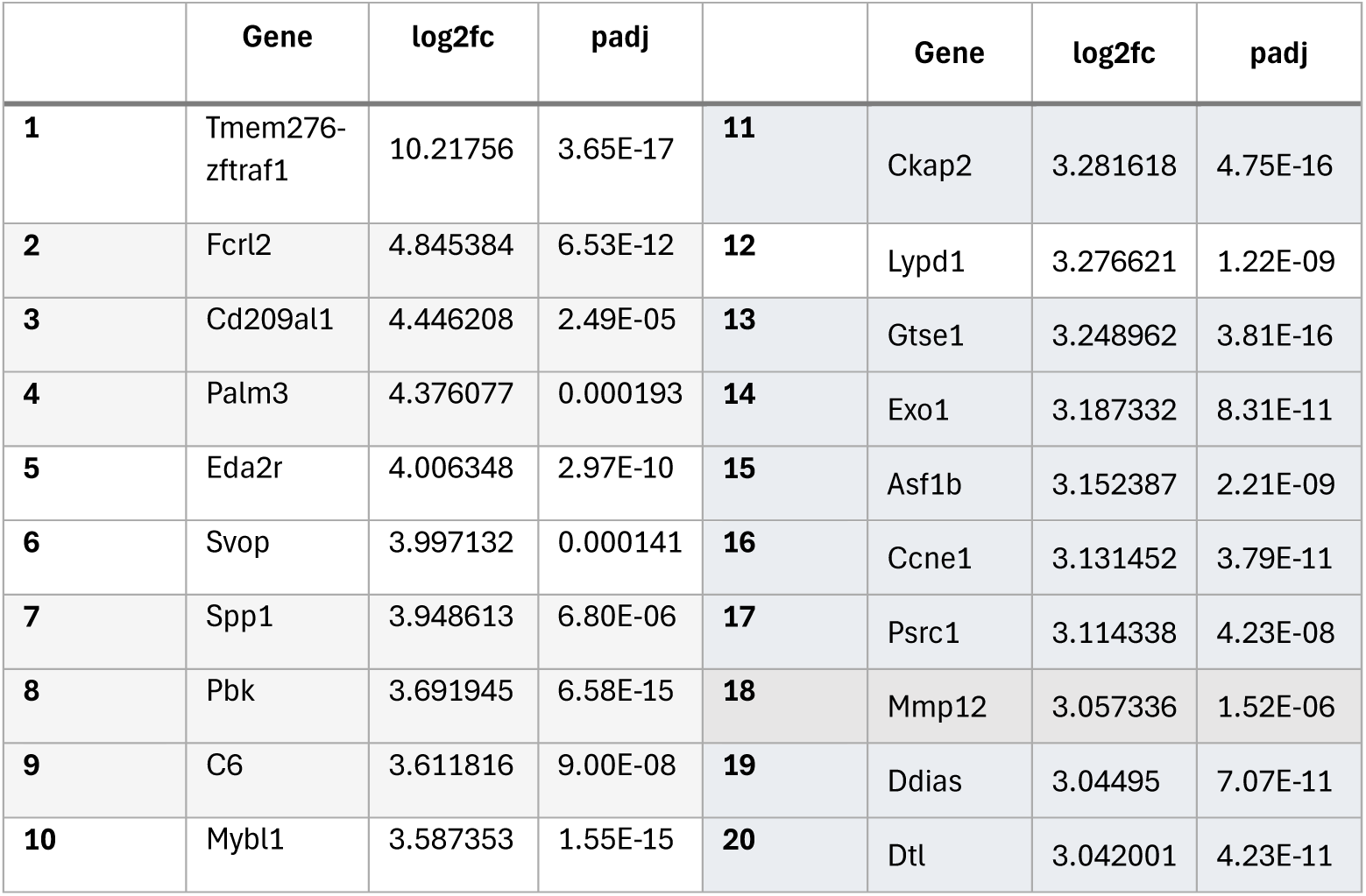
Top 20 differentially expressed genes in MCT/LPS compared with LPS treated rats. Genes involved in inflammation are shaded in grey and those in cell cycle and cell death in blue/grey.

KEGG biological pathway analysis demonstrated activation of pathways associated with cell division and DNA damage and repair (Falconi anaemia and p53 signaling pathway) in MCT-treated animals compared to DMSO (data not shown). LPS treatment stimulated cytokine responses and inflammatory pathways, as expected (Figure 5A). Similar changes in biological pathways were seen in DMSO compared to MCT/LPS to those seen with LPS alone, but additional biological pathways were induced such as the complement and coagulation cascades in MCT/LPS lung (Figure 5A). KEGG pathway analysis for LPS compared to MCT/LPS again suggested dysregulation of the cell cycle but additionally identified increased transcripts associated with neutrophil extracellular trap (NET) formation in MCT/LPS samples (Figure 5A). Gene ontology pathway analysis was also performed with similar outcomes: association of MCT treatment with induction of gene sets involved in cell division (data not shown); inflammatory responses, including the acute phase response and leukocyte migration and recruitment in LPS treated animals; and differences between LPS and MCT/LPS stimulated gene sets largely associated with cell division (Figure 5B). Interestingly, and in contrast to LPS, the MCT/LPS treated response identified upregulation of gene sets associated with Th1 immune responses, compared to DMSO alone (Figure 5B).

**Figure 5.**
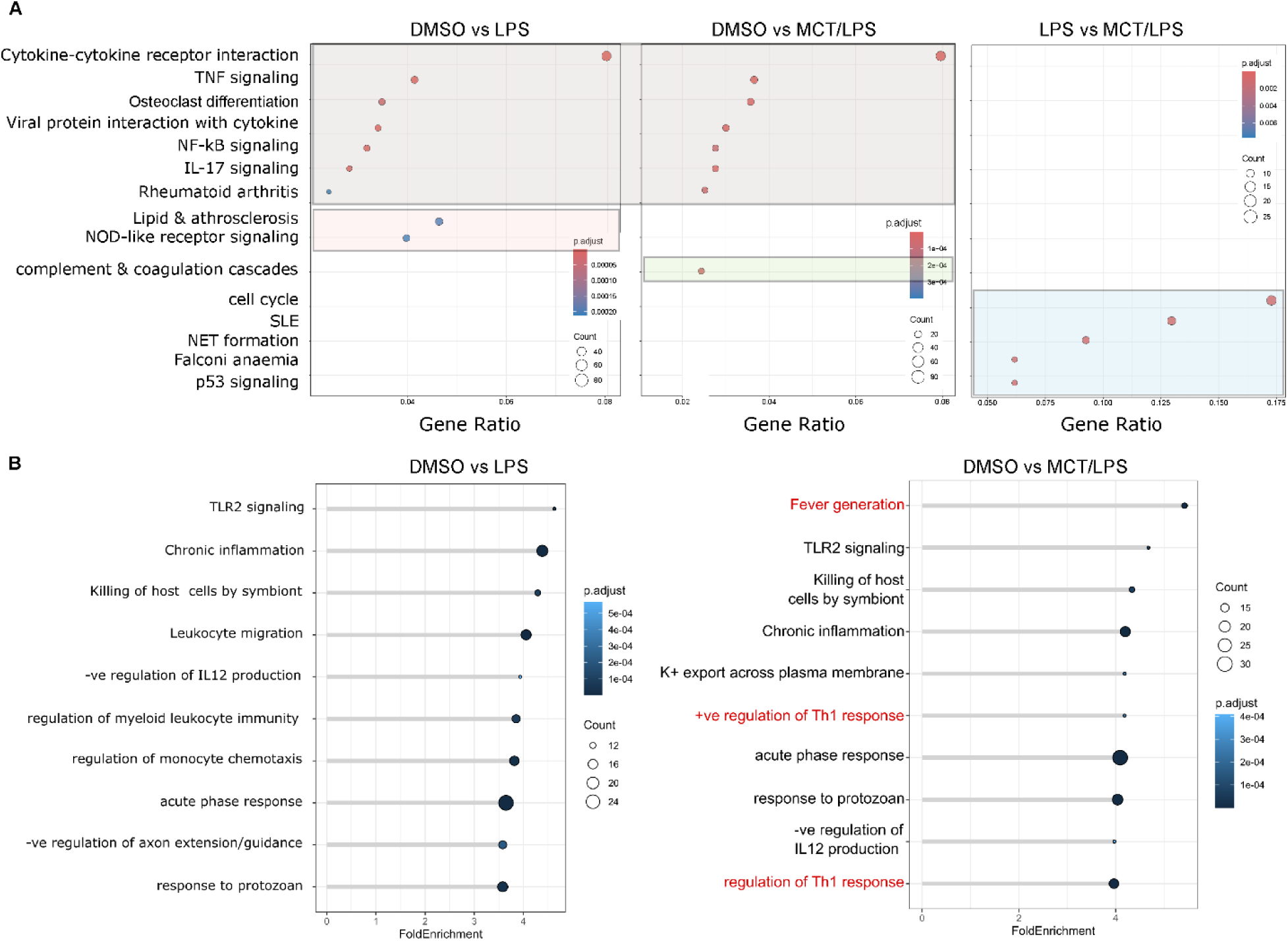
Pathway analysis highlights differences in the MCT, LPS and MCT/LPS treated rat lung. **A.** KEGG pathway analysis. Grey box=comparable pathways identified in DMSO compared to LPS or MCT/LPS, pink box =pathways specific to DMSO vs LPS, green=complement and coagulation pathway altered specifically in DMSO vs MCT/LPS and blue=pathways different LPS compared to MCT/LPS; **B.** Gene Ontology analyses. Gene sets enriched in DMSO vs MCT/LPS but not DMSO vs LPS are highlighted in red.

Next the changes quantitated by RT-PCR were interrogated in the RNAseq data, showing similar outcomes, including the striking induction of IFN-ϒ in MCT/LPS compared to LPS alone (data not shown). There was a significant correlation between changes detected by PCR with RNAseq for LPS (R^2^=0.9069) and MCT/LPS (R^2^=0.9633). This prompted analysis of the RNAseq data for mRNA changes associated with Th1/M1 phenotype, noting an M1 phenotype gene set was not part of the original gene ontology pathway dataset. Comparison of the normalised counts from RNAseq data demonstrated either no change or a decline in CD3, CD4 or CD8 mRNA (Figure 6A) but significant increases in the macrophage markers CD80 and CD86 specifically in the MCT/LPS but not LPS alone treated tissue (Figure 6B). PtprC, encoding the leukocyte marker CD45 was upregulated in LPS or MCT/LPS treated lungs compared to DMSO, but interestingly down regulated in MCT/LPS compared to LPS alone (Figure 6A). While mRNA for the cytokines IL6 and CCL9 were upregulated in LPS and MCT/LPS treated lungs, the mRNAs for CCL2, IL12b and Trem2 were further significantly upregulated in MCT/LPS compared to LPS alone (Figure 6C). Normalised enrichments scores (NES) for a set of M1-associated genes (Il6, Il12b, Ccl9, Cd80, Ccl22, Cd86) suggest enrichment of M1 phenotype with MCT treatment alone compared to DMSO (p=0.0093; NES=1.841), a decrease in enrichment with LPS treatment (p=0.0199; NES=1.588) and a highly significant and greater normalised enrichment with combined MCT/LPS treatment (p=3.02×10^-5^; NES=1.996).

**Figure 6.**
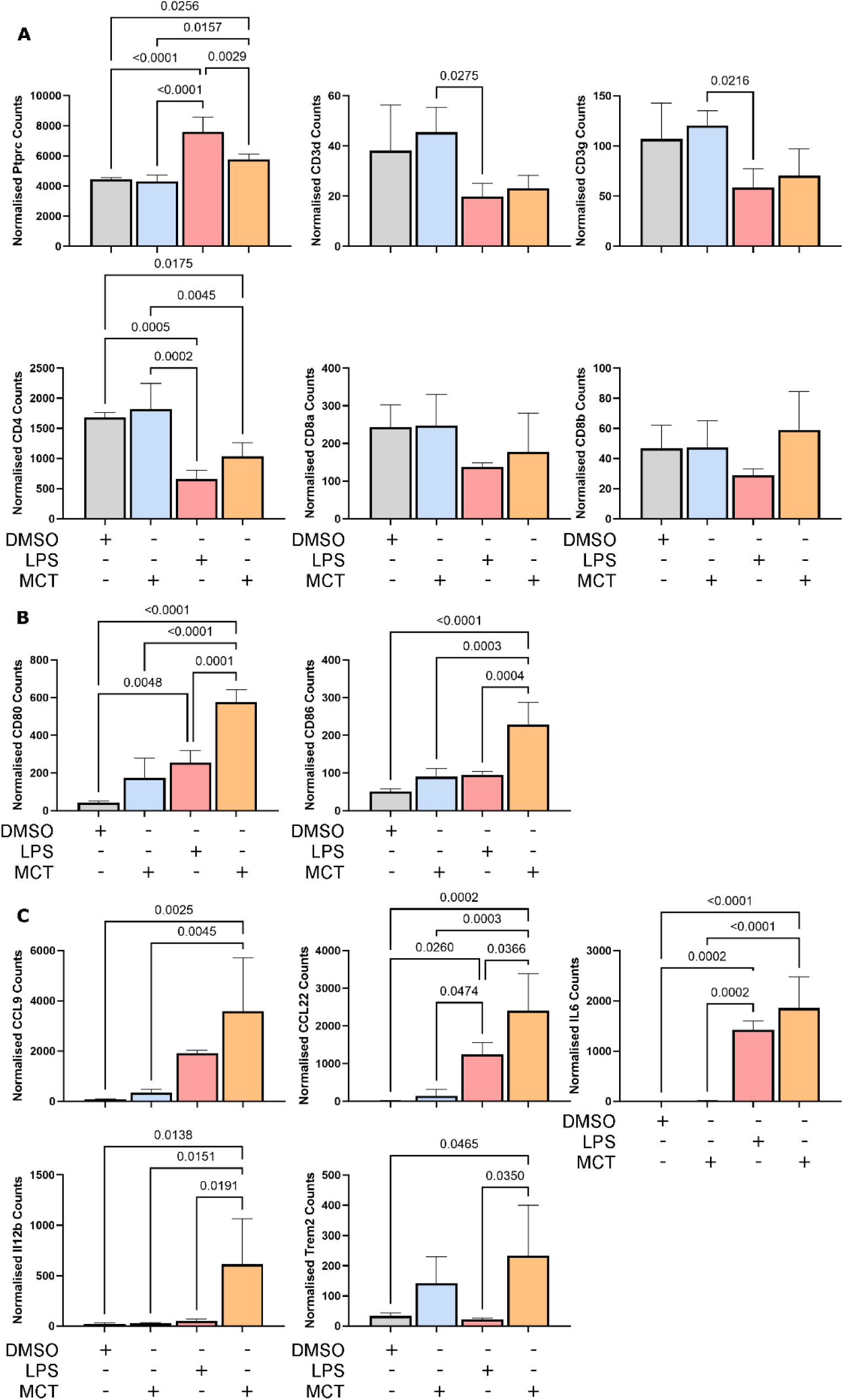
RNAseq analysis suggests induction of an M1 phenotype in MCT/LPS treated rat lungs. Rats were treated with DMSO or 60mg/kg MCT and at 14 days post treatment vehicle or LPS delivered intratracheally. 24 h later lungs were collected, total RNA extracted and subjected to RNAseq. Data was analysed for specific RNA levels for genes known to be associated with an M1/Th1 phenotype. **A.** Prtcp (CD45) and T-cell markers; **B.** Macrophage markers; C. M1/Th1 cytokine. RNAseq counts were normalised using DEseq’s “Median of Ratios” method and represent mean SEM, n=4 per group. Data were analysed by One-way ANOVA with Tukeys multiple comparisons, p values for significant difference are shown.

Combined, DGE and pathway analysis suggests important changes in the MCT/LPS treated rat lung that would be expected to promote inflammation, thrombosis and fibrosis (Figure 7), with increased complement activity (Figure 7A); dysregulation of the renin-angiotensin system (Figure 7B); increased fibrinogen and A2M (Figure 7C) and an overall alteration of the microenvironment in the alveoli with increased iron, increased M1 responses and increased NET formation (Figure 7D).

**Figure 7.**
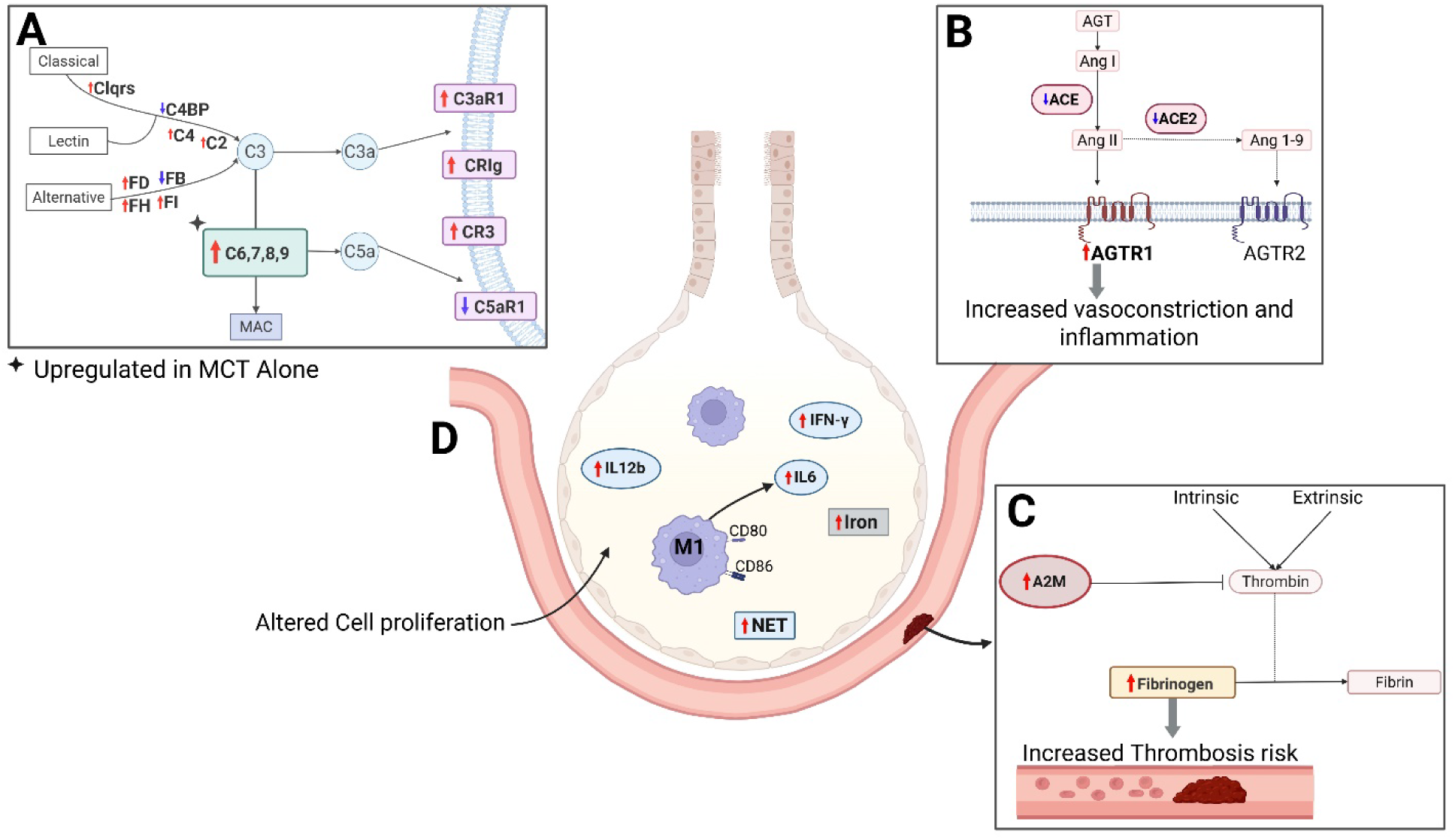
Schematic summary of pathways suggested to be altered in the MCT/LPS rat lung. **A.** complement pathway, increases in the classical and lectin pathways (C1qrs, C4 and C2, decreased C4binding protein (C4BP)), the alternative pathway (increased factor D, factor H and factor I but decreased factor B), terminal complement components that promote the membrane attack complex (MAC) (C6, 7, 8 and 9), and increased complement C3a receptors (C3aR1, CrIg and CR3) but decreased C5a receptor (C5aR1). Together this would promote C3a-driven inflammation and cell lysis by the MAC; **B.** Renin-angiotensin system, with down regulation of ACE and ACE2 but upregulation of AGTR1, predicted to promote vasodilation and inflammation; **C.** coagulation pathway, with increased alpha 2 macroglobulin (A2M) that would inhibit thrombin cleavage of fibrinogen, and increased fibrinogen itself that would promote the risk of thrombotic events; **D.** Predicted lung alveolar environment with **increased iron levels** (eg. increased ERFE that would inhibit hepcidin increasing systemic iron mobilization; increased **NET formation** (eg. gene expression such as citt13, HMGB1 and NE but decreased LL37 and PAD4), an **M1 phenotype** (increased CD80, CD86, IL6, IL12b and IFN-ϒ), and **altered cell proliferation** (eg. increased mRNA for DNA damage such as Gtse1 and Dtl1 and growth and differentiation, such as pannexin 3). Supporting information for specific DGE changes are available in supplementary data file. Image generated with BioRender.

## DISCUSSION

The COVID-19 pandemic has highlighted a need for flexible laboratory models of inflammatory lung disease and ARDS. Our prior review proposed that the MCT-treated rat lung reflects many key pathologies seen in the COVID-19 lung (Kris et al. 2025). Our experimental assessment here demonstrates that MCT treatment alone does not have a major impact on respiratory function and does not meet the criteria of an ALI model (Kulkarni et al. 2022). Pretreatment of rats with MCT, however, does predispose to worse respiratory and altered inflammatory outcomes following stimulation with a model pathogen, LPS, 14 days later and some of these key changes are consistent with the COVID-19 lung and ARDS.

Treatment of adult rats with a single dose of MCT (50-100 mg/kg) or the active metabolite MCTP leads to PH at 28 days post treatment (Gillespie et al. 1985; Gomez-Arroyo et al. 2012; Hill, Gillespie, and McMurtry 2017). The MCT PH model is progressive and earlier, at 14 days post treatment, animals are pre-hypertensive with indicators of inflammation and pulmonary vascular dysfunction (Lai, Olson, and Gillespie 1991). The study here defined that a single low dose (15 mg/kg) or shorter time frame post MCT treatment (3 or 7 days), compared to higher dose (60 mg/kg) and 14 days post MCT treatment of an aged rat (approximately 12 month) did not show measurable impact. One animal, in the MCT-treated group did show unusual gross pathology of the lung and suggestion of a haematoma of the liver, but it is not clear if these were age associated or resulting from the MCT treatment. These changes were seen at termination of the experiment and were not fatal.

Subsequent focus on the 60 mg/kg MCT treatment 14 days prior to low dose, 24 h LPS stimulation of the rat lung did induce different responses, with a decrease in lung compliance and increase in elastance and resistance with MCT/LPS treatment. Both wet and dry weight was increased, suggesting both oedema and increased cellular or fibrotic tissue mass. The increase observed in total protein content in the BAL supports the increased production of an exudate and/or a loss of alveolar barrier integrity. Pathology scoring of alveolar thickening and cellular infiltrate demonstrated these were increased in MCT/LPS treated lungs, but not significantly above that seen with LPS alone. Staining of the BAL fluid demonstrated that in both instances there was a substantial lymphocytic cell population with large macrophage-like cells likely representing alveolar or activated macrophages that were particularly prominent in the MCT/LPS rat lung. These large macrophage-like cells are similar to those described in the BAL of animals with other lung insults such as in surfactant protein-D deficient mice (Botas et al. 1998), although a more detailed identity of these cells, for instance as recruited or tissue resident macrophages (Aegerter, Lambrecht, and Jakubzick 2022) remains to be determined. Together, these MCT/LPS-induced changes meet three of the four domains suggested to reflect the criteria for a model of ALI: histological damage; alveolar-capillary barrier changes and lung dysfunction (Kulkarni et al. 2022).

The fourth domain of inflammation was demonstrated, by RT-PCR with induction of a proinflammatory response (TNF, C3 and CXCL10) by LPS that were not significantly altered in MCT/LPS treatment. Strikingly, MCT/LPS but not LPS alone induced mRNA for IFN-ϒ and this set the rationale for a more detailed analysis of the transcriptional changes in the MCT model by RNAseq. Principle component analysis from RNAseq data supported that there were not major changes in the MCT compared to DMSO control treated rat lungs. There were, however, significant upregulation at a molecular level of pathways involved in inflammation, cell division/cell death and components such as erythroferrone (ERFE) and lipocalin-2 (LCN2). ERFE binds to ligands of the bone morphogenic protein (BMP) receptor and indirectly inhibits the expression of hepcidin, a molecule responsible for mobilising iron stores (Coffey and Ganz 2018). Iron homeostasis and accumulation in lung macrophages is associated with lung dysfunction and damage (Neves et al. 2017). Rats provided an iron restrictive diet can resolve the MCT-induced right ventricular failure, suggesting high iron is involved in mediating at least part of the MCT-induced pathology (Naito et al. 2013) and MCT has also been linked to altered BMP signaling (Morty et al. 2007). Additionally, LCN2 has been associated with inflammation and fibrosis in the lung (Galaris et al. 2023) and is induced in response to LPS and promotes macrophage recruitment and inflammation, including IL-6 production, for instance through increased iron levels in macrophages in the lungs of mice or cells *in vitro* (An et al. 2023). Overall, changes in ERFE and LCN2 in the MCT-treated are consistent with prior literature on MCT and LPS responses and would be expected to disrupt iron homeostasis and inflammation (Figure 7).

In contrast to these smaller changes induced by MCT alone, the transcriptome of MCT/LPS treated animals clearly segregated from DMSO or MCT as well as LPS treated alone, confirming that the MCT pre-treatment predisposes to an altered response to LPS. Comparison of the top 20 up-regulated mRNAs demonstrated many similar changes compared to LPS, for instance IL6, Saa3, CXCL11 and NOS2 induction. An inflammatory cytokine response, and in particular IL6, has been a key factor identified and point of treatment for the COVID-19 lung (Blanco-Melo et al. 2020; Derde et al. 2025). Chemokines such as CXCL11 and reactive oxygen species/oxidative stress are established inflammatory mediators of respiratory pathology and ARDS. For instance, antagonism of CXCR7, a receptor for CXCL11, leads to reduced LPS induction of vascular permeability and neutrophil infiltration in mice (Pouzol et al. 2021). The MCT/LPS stimulated lung had increased mRNA for further genes such as pannexin (Panx) 3. Panx are ion channels that can regulate cell proliferation, with a growing interest in immunomodulatory roles (Santiago-Carvalho, Ishikawa, and Borges da Silva 2024) and thus Panx3 may be a factor of future interest in lung disease. Direct comparison of the highest differentially expressed genes in MCT/LPS compared to LPS alone, highlighted multiple inflammatory-related genes such as CD209 (DC-SIGN), paralemmin 3 (Palm3) and Spp1. Both Palm3 and Spp1 have been related to alveolar macrophage inflammation (Chen et al. 2017), with an Spp1 transcriptional signature in human alveolar macrophages associated with pulmonary fibrosis (Reyfman et al. 2019).

C6, a part of the complement membrane attack complex (MAC) was a highly upregulated mRNA in MCT/LPS rat lung, with the complement pathway identified by KEGG analysis as upregulated in MCT/LPS rat lung. Pathway analysis identified other components (C1q, C2, C3 and C4), receptors (C3aR1, CrIg) and regulators (FH, FD) of the complement pathway as altered in the MCT/LPS model. The complement system is involved in various lung disease states (Kulkarni et al. 2018; Pandya and Wilkes 2014), and an atlas of complement producing cells in the lung, derived from single cell RNAseq data (Chaudhary et al. 2022), suggests multiple cell types of the lung likely contributing to these complement changes, such as C2, C3, C4 and FH from mesothelial cells and C1q from macrophages. Prior characterization of complement in the lung has strengthened the relevance of the alveolar macrophages, for instance in C2 production (Ackerman et al. 1978) or CrIg-mediated phagocytosis (Nagre et al. 2018) suggesting both the complement system and macrophages could be important in the MCT/LPS response in the rat lung (Figure 7A, D).

MMP12 was upregulated by MCT treatment alone and in MCT/LPS treatment compared to LPS. MMP12 is a matrix metalloproteinase produced by alveolar macrophages that is involved in tissue remodeling (Shapiro, Kobayashi, and Ley 1993). The enzyme targets elastin and extraceullar matrix proteins such as fibronectin, type IV collagen and glycosaminoglycans such as heparan sulfate (Gronski et al. 1997) and MMP12 is a driver of Fas-induced pulmonary fibrosis in mice (Matute-Bello et al. 2007). MMP12 activity is increased by iron and would be expected to have exacerbated proinflammatory actions in the context of high iron, as described above due to increased Erfe and Lcn2, in the MCT-treated rat lung.

These transcriptional changes in the MCT treated aged rat alone or MCT/LPS compared to LPS suggest benefit for using this model to study the roles of selected factors, such as Panx3, Palm3, Spp1, the complement system or MMP12, in ALI or pathogen responses in the lung. Both RT-PCR and RNAseq analysis demonstrated that LPS stimulation decreased mRNA levels for ACE2. Reduced cellular ACE2 would be expected to lead to increased AngII, that would be predicted to lead to inflammation and vasoconstriction, as is associated with ARDS (Hrenak and Simko 2020; Imai, Kuba, and Penninger 2006; Marshall et al. 2002). Reduced ACE2 has been previously described in MCT-associated PH in the rat (Malikova et al. 2016) and in the COVID-19 lung (Pagliaro and Penna 2020), potentially by receptor internalization and degradation (Lu et al. 2022), or enzymatic cleaved from the cell surface during the inflammatory response, such as by actions of ADAM-17 (Sun et al. 2024). In contrast, ELISA quantitation demonstrated increased ACE2 in the BAL fluid of LPS stimulated lungs and a further increase in MCT/LPS treatment, which correlates with increased soluble ACE2 in COVID-19 patients (Gerard et al. 2021). This was accompanied by increased protein levels in the BAL, that may suggest lung injury. Pathway analysis suggests that the impact of MCT on the RAS is not just via ACE2, but also a decrease in ACE and an increase in AGTR1. Together dysregulation of the RAS would contribute to a pro-inflammatory, thrombotic and fibrotic environment (Figure 7B) (Gaddam, Chambers, and Bhatia 2014).

Analysis of the RNAseq data for our mRNAs previously defined by RT-PCR, confirmed the significant and unique upregulation of IFN-ϒ in MCT/LPS compared to LPS alone. The cytokine environment was specifically enriched for mRNAs known to be associated with an M1 phenotype, such as IL6, CCL9, CCL22 and IL12b in MCT/LPS compared to LPS alone treated lungs. The macrophage lineage was seen to be associated with this response with no induction of T-cell markers, CD3, CD4 or CD8, but significant induction of CD80 and CD86 in MCT/LPS compared to LPS alone. These specific mRNA changes are consistent with the identification Th1 response pathways by GO, since an M1 profile is not available in this dataset. Data is also consistent with reported increased M1/M2 ratio measured by flow cytometry in lung tissue and alveolar lavage fluid 6 days after 50mg/kg MCT treatment of rats (Tang et al. 2021) (Figure 7D). In contrast, triggering receptor expressed on myeloid cells-2 (TREM-2) was significantly upregulated in MCT/LPS compared to LPS alone. TREM-2 is a factor aligned with M2 responses in the lung, with down regulation in LPS macrophages and a murine model of ALI, and a protective role in sepsis-induced ALI (Shen, He, and Lin 2025).

In addition to the major gene expression changes discussed above, GO or KEGG pathway analysis suggested upregulation of pathways for coagulation, NET formation and cell death. Cell death pathways were upregulated in MCT alone and MCT/LPS compared to LPS alone for p53 signaling and Falconi anaemia, reflecting changes in mRNAs involved in the cell cycle, cell division and cell death. The induction of gene expression profiles associated with cell cycle regulation seen following MCT treatment is also consistent with prior literature, such as increased cell proliferation seen during development of MCT-induced PH. The coagulation pathway was specifically upregulated in MCT/LPS rat lungs but not LPS alone, with upregulation of alpha 2 macroglobulin (A2M), which is known to inhibit thrombin. Co-incident with upregulation of fibrinogen induced by MCT/LPS this would be expected to increase propensity to clot and a thrombotic state (Figure 7C). One last key gene set identified by GO analysis as increased in MCT/LPS compared to LPS treated lungs is neutrophil extracellular trap (NET) formation (Zhou et al. 2024) (Figure 7D). Although our analysis suggests importance of macrophages, neutrophils are well described as responders in ALI and associate with changes such as increased MMP, IL-6 and TNF and alveolar inflammation in ARDS (Bos and Ware 2022). The process of NETosis – production of an extracellular matrix to trap neutrophils and pathogens, is important in multiple lung pathologies including non-infectious disease (Jorch and Kubes 2017) and in the progression of PH in MCT-treated rats (Semenkova et al. 2025). The addition of the MCT prestimulation in this model, thus enhances these important known mechanisms of lung inflammatory pathology and would benefit the study of these processes.

The implications of our findings are that this model of MCT pretreatment reflects an aged rat with developing PH and potential altered lung responses to pathogen stimuli (such as LPS), that are compounded by cardiovascular and pulmonary settings, likely reflective of the complexity of human ARDS patients. The data here suggests that prior MCT exposure itself does not cause an ALI but establishes a subclinical change in the lung that has detectable molecular changes associated with inflammation and cell proliferation, that will predispose the lung to respond to LPS towards an M1 inflammatory environment. This is associated with transcriptional changes in key systems such as alveolar macrophages, the RAS, increased iron, complement and coagulation pathways and NET formation, that we know are associated with lung damage and disease. It may be useful to incorporate the complexity and inflammatory bias this model provides to aid in understanding the relationships between multiple systems that are part of the response to pathogens and can influence patient outcomes in clinical settings such as ARDS and the COVID-19 lung.

## ACKNOWLEDGMENTS

Thank-you to the College of Medicine and Public Health pre-clinical services team for care of animals used in this study, and Flinders Microscopy and Microanalysis for use of equipment for tissue processing and microscopy.

## AUTHOR CONTRIBUTIONS

Conceived and designed research (JMC, SB, DLD), performed experiments (DDB, EC, TP, LK, JMM), analyzed data (DDB, EC, TP, LK, MR), interpreted results of experiments (JMC, SB, DLD, JMM), prepared figures (JMC, DDB, EC, TP) drafted manuscript (JMC), edited and revised manuscript (JMC, SB, DLD, DDB, TP, EC, LK, MR), approved final version of manuscript (JMC, SB, DLD, DDB, TP, EC, LK, MR).

