## Supplemental Table 1 for "Monocrotaline treatment of the rat predisposes to altered lipopolysaccharide-induced lung responses"

**Supplementary Table 1. Primers used for RT-PCR.** All primers are designed against *Rattus norvegicus* mRNA sequences

| **Target mRNA** | **Forward Primer** | **Reverse Primer** | **Amplicon length (bp)** | **Accession Number** |
| --- | --- | --- | --- | --- |
| **ACE2** | TCGGAGTTCATAGTGCCACG | CCAGCAGGAGCTTGACATCT | 244 | XM_039099655.1 |
| **CCL5** | CTCCCTGCTGCTTTGCCTAC | ACTTGGCGGTTCCTTCGAG | 124 | NM_031116.3 |
| **CTSL** | CGACACAGGGTTTGTGGATA | ATCGCTGTCCATCAATTCAC | 354 | NM_013156.2 |
| **CXCL10** | ATGAACCCAAGTGCTGCTGT | CTCTCTGCTGTCCATCGGTC | 347 | NM_139089.2 |
| **GAPDH** | GGCAAAAGGACGGTAACACG | GTGCTGGATCTGTGGGTTGT | 215 | NM_138880.3 |
| **IFNγ** | GGCAAAAGGACGGTAACACG | GTGCTGGATCTGTGGGTTGT | 215 | NM_138880.3 |
| **IL2** | TCACTTGGAAGACGCTGGAA | TGGCTCATCATCGAATTGGC | 103 | NM_053836.1 |
| **IL6** | CACTTCACAAGTCGGAGGCT | AGCACACTAGGTTTGCCGAG | 502 | NM_012589.2 |
| **IL17a** | GTGAAGGCAGCGGTACTCAT | ATGTGGTGGTCCAACTTCCC | 272 | NM_001106897.1 |
| **NRP1** | TCATTCAAGGTGGGAAGCAC | GTGGCTCTCTCGGGGTAGAT | 214 | NM 145098.2 |
| **PPIA** | CTTCGACATCACGGCTGATGG | CAGGACCTGTATGCTTCAGG | 266 | NM 017101.1 |
| **TMPRSS2** | CACCTGCCATCCACATACAG | CCAGAACTTCCAAAGCAAGC | 140 | NM 130424.3 |
| **TMPRSS11a** | TGGGGAGTCCCAAAATGAGC | CCATGGCGAGTTAGAGACCC | 317 | XM_039092583.1 |
| **TNFα** | ACTGAACTTCGGGGTGATCG | GCTTGGTGGTTTGCTACGAC | 153 | NM_012675.3 |
| **Viperin** | ACTATTTGGACATTCTTGCTATCTC | AATCAGGAGGCATTGGAAA | 248 | NM_138881.2 |
