## Supplemental Figures for "Monocrotaline treatment of the rat predisposes to altered lipopolysaccharide-induced lung responses"

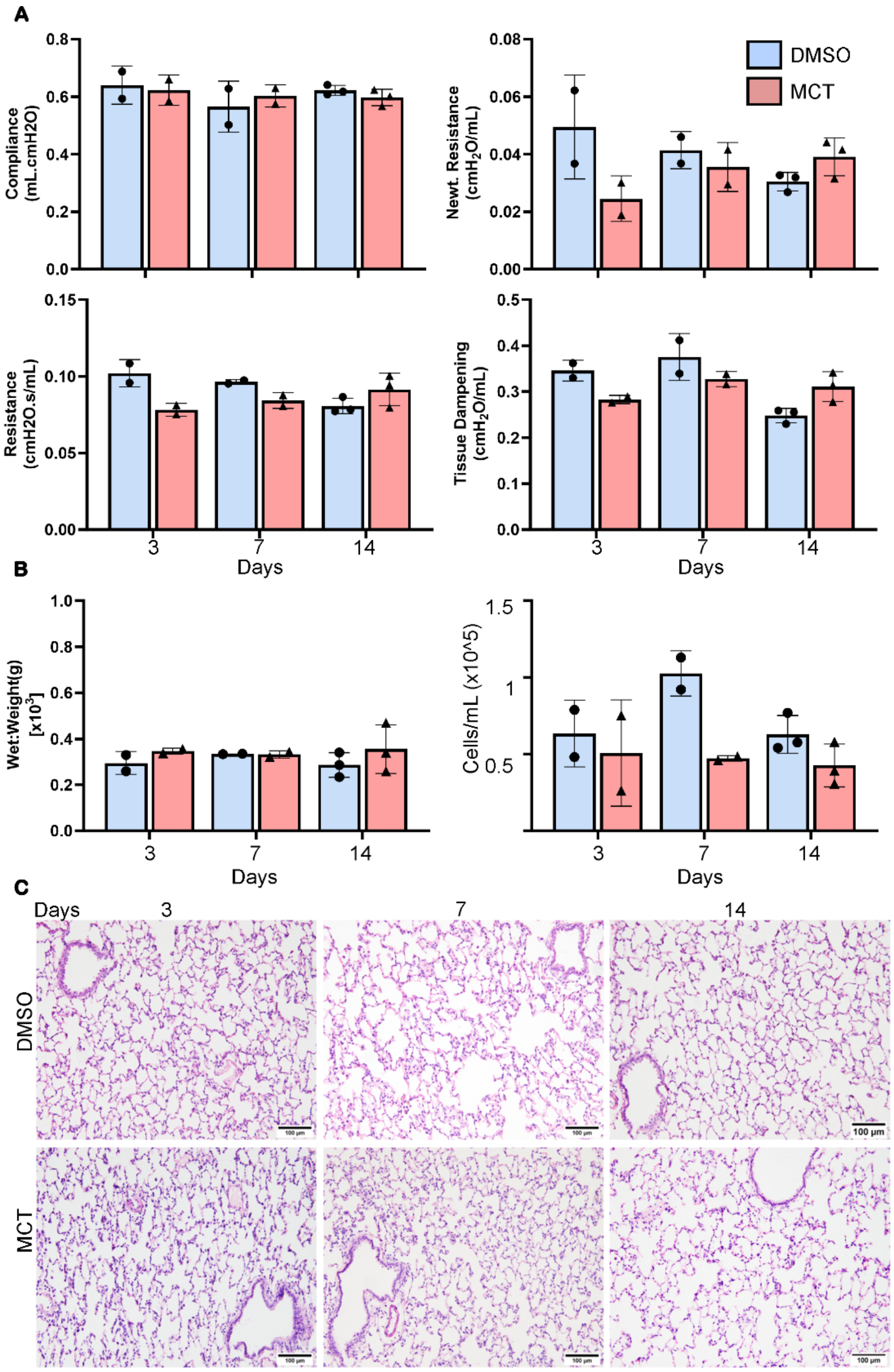


**Supplemental Figure 1. Time course and impact of MCT dose on the rat lung.** Rats were treated with DMSO or MCT, n=2 per group at each time point. At the time point indicated **A.** respiratory mechanics were analysed; **B.** lungs were removed and wet weight determined. BAL was extracted and cells counted by light microscopy; Individual data points represent individual animals. **C.** lungs were fixed and processed for H&E staining with images collected by bright field microscopy. Representative images are shown.

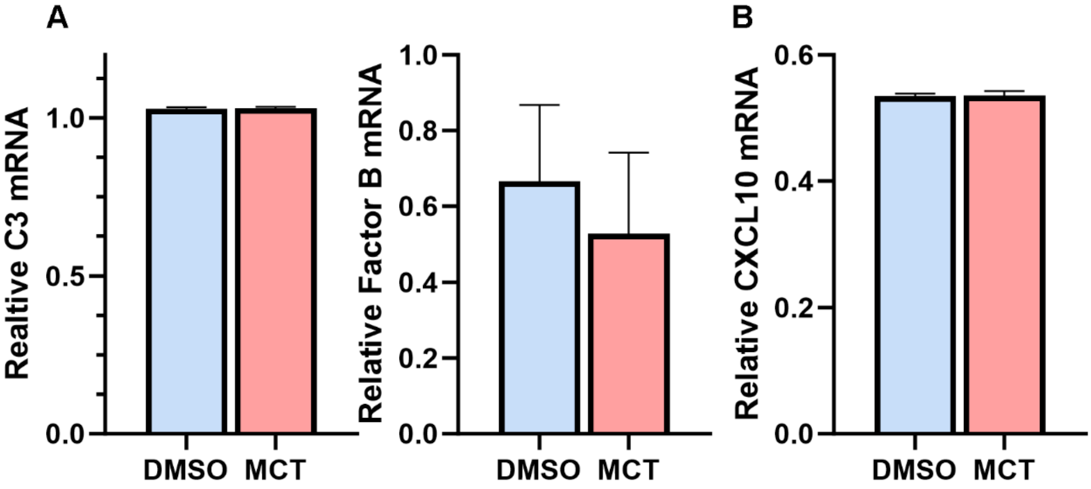


**Figure 2. MCT has no impact on liver mRNA levels.** Rats were treated as in Figure 1 and liver collected at 14 days post 60 mg/kg MCT or DMSO treatment. Total RNA was extracted and subjected to RT-PCR for **A.** complement components C3 and complement factor B; **B.** the inflammatory cytokine CXCL10. Results were normalised against cyclophilin and represent n=2 per group. SD values are shown.
